# Therapy-induced transdifferentiation promotes glioma growth independent of EGFR signaling

**DOI:** 10.1101/2020.06.02.130948

**Authors:** Hwanhee Oh, Inah Hwang, Lingxiang Wu, Dongqing Cao, Jun Yao, Haoqiang Ying, Jian Yi Li, Yu Yao, Baoli Hu, Qianghu Wang, Hongwu Zheng, Jihye Paik

**Affiliations:** Department of Pathology and Laboratory medicine, Weill Cornell Medicine, New York, NY 10065, USA; Meyer Cancer Center, Weill Cornell Medicine, New York, NY 10065, USA; Department of Bioinformatics, Nanjing Medical University, 211166, China; Neurosurgical Immunology Laboratory, Neurosurgical Institute of Fudan University; Department of Neurosurgery, Huashan Hospital, Fudan University, Shanghai, China; Department of Molecular and Cellular Oncology, University of Texas MD Anderson Cancer Center, Houston, TX 77054, USA; Department of Pathology and Lab Medicine, North Shore University Hospital and Long Island Jewish Medical Center, Northwell Health, Lake Success, Donald and Barbara Zucker School of Medicine at Hofstra/Northwell, NY 11042, USA; Children’s Hospital of Pittsburgh Department of Neurological Surgery, University of Pittsburgh School of Medicine, PA 15224, USA

**Author notes:** These authors contributed equally (Hwanhee Oh, Inah Hwang). **Author Contributions**: Conception and design, H.O. H.Z., J.P.; Development of methodology, H.O., B.H., Q.W.; Acquisition of data, H.O., I.H., Y.Y., D.G.; Analysis and interpretation of data, H.O., I.H., J.Y., Q.W., B.H., Y.Y. H.Z., J.P.; Writing, review and/or revision of the manuscript, H.O., I.H., Y.Y., H.Z., J.P.; Administrative, technical, or material support, Y.Y., J.Y.L., H.Z. J.P.; Study supervision, Y.Y., Q.W., H.Z., J.P.

**Keywords:** EGFR, Slug, YAP, glioma, therapy resistance, lineage transdifferentiation

## Abstract

Epidermal growth factor receptor (EGFR) is frequently amplified, mutated and overexpressed in malignant gliomas. Yet the EGFR-targeted therapies have thus far produced only marginal clinical response, and the underlying mechanism remains poorly understood. Through analyses of an inducible oncogenic EGFR-driven glioma mouse model system, our current study reveals a small population of glioma cells that can evade therapy-initiated apoptosis and potentiate relapse development by adopting a mesenchymal-like phenotypic state that no longer depends on oncogenic EGFR signaling. Transcriptome analyses of proximal and distal treatment responses further identify TGFβ/YAP/Slug signaling cascade activation as major regulatory mechanism that promotes therapy-induced glioma mesenchymal lineage transdifferentiation. Following anti-EGFR treatment, the TGFβ secreted from the stressed glioma cells acts to promote YAP nuclear translocation and activation, which subsequently stimulates upregulation of the pro-mesenchymal transcriptional factor Slug and then glioma lineage transdifferentiation towards a stable therapy-refractory state. Blockade of this adaptive response through enforced dominant negative YAP expression significantly delayed anti-EGFR relapse and significantly prolonged animal survival. Together, our findings shed new insight into EGFR-targeted therapy resistance and suggest that combinatorial therapies of targeting both EGFR and mechanisms underlying glioma lineage transdifferentiation could ultimately lead to deeper and more durable responses.

**Significance:** This study demonstrates that molecular reprogramming and lineage transdifferentiation underlie anti-EGFR therapy resistance and is clinically relevant to the development of new combinatorial targeting strategies against malignant gliomas carrying aberrant EGFR signaling.

## Introduction

Malignant glioma is the most common and lethal type of primary brain tumor (1–3). In its most aggressive form, glioblastoma (GBM) has a median survival of only 12-15 months even after maximum surgical resection and chemoradiotherapy. The statistics have barely changed over the past twenty years (4,5). But contrary to the relative lag of clinical advancement, the world has witnessed an explosion of knowledge in glioma biology and basic science discovery over the same time period. Particularly, the breakthrough in sequencing technology has finally made unraveling the complete genomic landscape of GBMs a reality (6–8). Among the complex genetic and genomic events, EGFR has attracted arguably the most attention due to the fact that its amplification together with mutation and/or rearrangement were identified in ~ 60% of GBM patient samples (7,9). The success of EGFR tyrosine kinase inhibitors (TKIs) in treatment of non-small cell lung carcinoma (NSCLC) patients carrying active EGFR mutations further made it an appealing target for therapeutic interventions in GBMs (10–13).

Aberrant EGFR activation triggers pro-survival and proliferative signaling cascades in GBM. Despite its prevalence and the demonstrated role, EGFR-targeted interventions by strategies such as small molecule TKIs, antibodies and vaccines, have failed to achieve tangible clinical benefit, even in patients with EGFR amplification/mutations (14,15). A variety of resistance mechanisms have been proposed, such as incomplete target suppression (16,17), intratumoral heterogeneity (18,19), activation of downstream effectors in the same signaling pathway or engagement of alternative survival pathways (20–22). In line with those findings, the vast majority of current efforts have been focused on developing better drugs or drug combinations to more vigorously suppress EGFR and its downstream surrogates. However, the clinical outcome of deep EGFR suppression and potential resistance development afterwards have been poorly understood.

A major hurdle in deeper investigation into aftereffects of deep EGFR suppression is the dearth of relevant experimental model systems. In human GBMs, high-copy *EGFR* amplification occurs within extrachromosomal DNA known as double minutes (19,23,24). Therefore, the amplified *EGFR* copies are numerically unstable and often lost in cultured tumor cells (25,26). This intrinsic instability also poses as an immense barrier for *in vitro* genetic manipulation and thus necessitates alternative experimental systems to study EGFR functions and treatment responses.

In an attempt to address this challenge, we previously developed a glioma animal model driven by inducible expression of a truncated oncogenic EGFR mutant (EGFR*) deleted of exon 2–7 in EGFR extracellular domain (27). By utilizing this model system, we revealed that development of resistance to EGFR-targeted therapy against malignant glioma occurred through EGFR-dependent and -independent mechanisms. In this study, we set out to further exploit this experimental platform and probe for the molecular mechanism(s) underlying EGFR-targeted therapeutic response and resistance development.

## Results

### No overt genetic changes in iEIP relapse tumors following EGFR* suppression

By utilizing animals engineered with doxycycline (dox)-off oncogenic EGFR* transgene and conditional Ink4a/Arf and Pten alleles (*Nestin-CreER^T2^*; *Ink4a^L/L^*; *Pten^L/L^*; *hGFAP_tTA*; *tetO-EGFR*\*, designated iEIP), we previously demonstrated that sustained oncogenic EGFR signaling was required for maintenance of EGFR-driven glioma progression and suppression of oncogenic EGFR* transgene expression induces tumor regression (27). But despite of a robust initial response, the regressed tumors relapsed inevitably and eventually led to mortality. To investigate underlying mechanisms of the tumor recurrence, we grafted a group of immunocompromised *nu/nu* mice orthotopically with luciferase-expressing iEIP glioma cells. Bioluminescence imaging (BLI) revealed formation of significant tumor burdens by 5-7 weeks. Consistent with our previous study (27), extinction of oncogenic EGFR* transgene expression by doxycycline treatment elicited a deep tumor regression, with virtually no BLI-detectable tumor mass by 2 weeks following dox switch (Fig. 1A). After a period of relative indolence that lasted ~ 6 – 10 weeks, however, the animals started to show signs of relapse even under continued doxy treatment. Eventually, all of them succumbed to tumor progression ranging from 12 to 20 weeks (Fig. 1B). Immunohistochemistry (IHC) analysis of the relapse tumors found no evidence of reactivation of EGFR and/or its downstream Erk signaling reactivation (Fig. 1C), suggesting that the relapse was unlikely to be driven by compensatory EGFR signaling.

**Figure 1.**
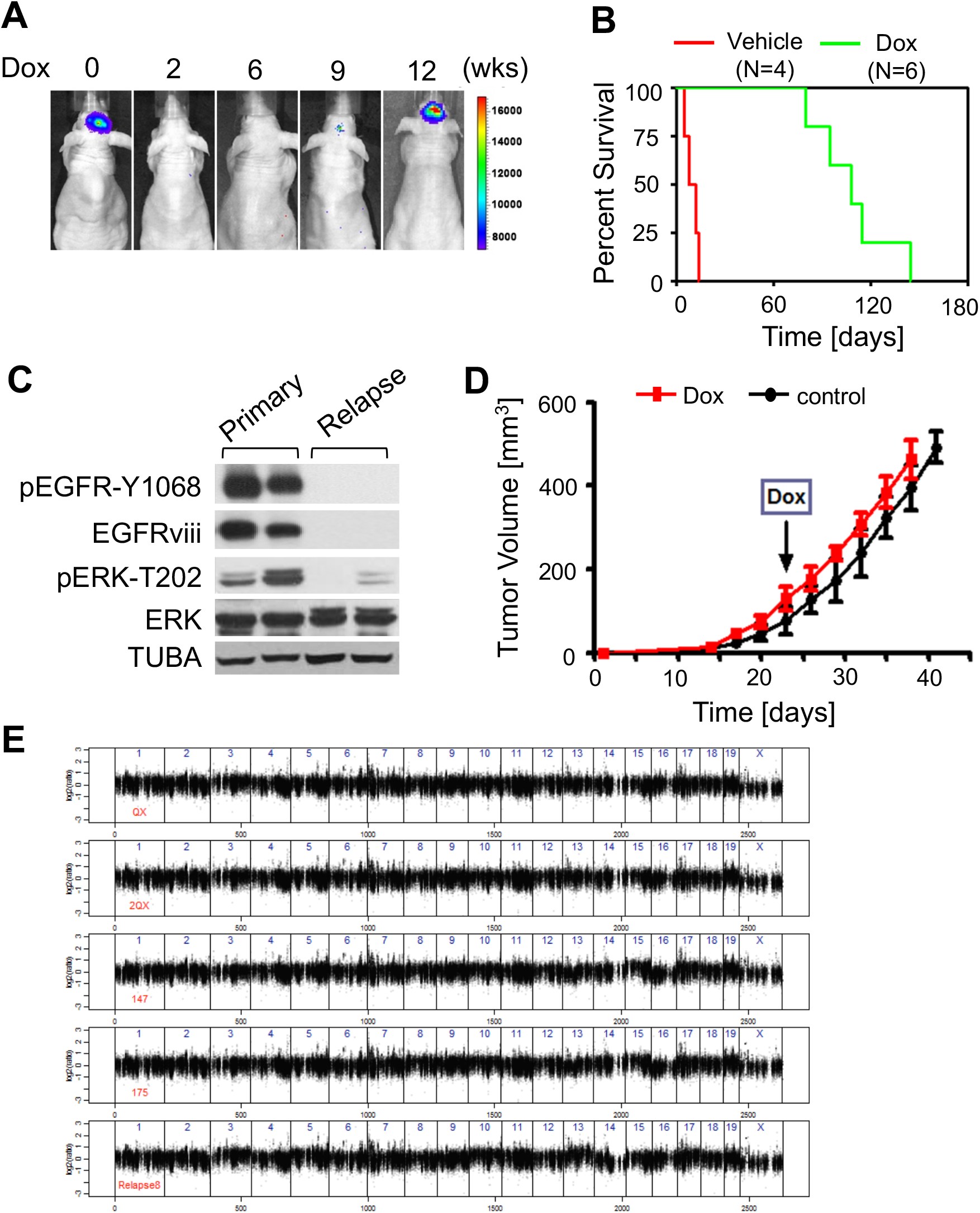
The relapsed tumor upon EGFR* inhibition is driven by oncogenic EGFR-independent pathway(s). **A**, Representative BLI on indicated days post dox treatment (N=5). **B,** Survival plot for mice treated with control (N=4) or dox water (N=6). **C,** Representative images of WB analysis for indicated proteins in primary and relapsed tumors. **D,** Relative tumor growth of re-grafted relapsed tumors after control or dox water treatment for the indicated time. Relapsed tumors from dox treatment (first relapsed) were re-grafted subcutaneously into *Nu/Nu* mice and maintained off-dox. After the tumors (second transplant) reached analyzable size (marked, ~200 mm^3^), the animals were separated into control (N=3) or dox (N=4) groups. **E,** WES copy number results for three relapse (147, 175, relapse8), and two matched parental (QX, 2QX) lines. Segmented copy number data is shown for each chromosome, by genomic position in columns and by cell lines in rows.

In targeted therapies a subpopulation of drug-tolerant cells are known to survive through potentially lethal exposures and drive relapse development. These therapy-tolerant states are often transient and can be reversed back to drug-sensitive state upon removal of the treatment. To assess whether the observed anti-EGFR resistance was a stable feature, freshly isolated iEIP relapsed tumor cells were re-transplanted subcutaneously into recipient nude mice. The animals were kept off-dox until tumors reached analyzable sizes. IHC analyses of the treatment-naïve controls revealed that only a small fraction (< 5%) of tumor cells in the re-grafted gliomas reactivated their EGFR* transgene expression (Fig. S1A). Moreover, re-initiation of dox-treatment exerted no visible effect on tumor growth (Fig. 1D), indicating that progression of the relapsed tumors no longer relies on oncogenic EGFR* signaling.

The finding that relapsed tumors had escaped oncogenic EGFR signaling addiction promoted us to search for potential genetic events that might fuel the resistance development. Three relapse and two matched treatment-naïve tumors derived from the same parental line were analyzed by whole exome sequencing (WES) (Fig. S1B). First analysis of copy number variations recovered no recurrent regions of copy number alterations (Fig. 1E). To identify the potential relapse-specific mutations, we first excluded the SNPs that appeared in both parental and relapse tumor cells. The remaining sequence data were then filtered to subtract the variants present in less than 25% of the reads. The rare variants that eluded the threshold filter and appeared only in relapse tumors affect 15 genes. However, further verification analysis of the raw data using integrated genomic viewer revealed that all of the 15 SNPs preexisted in the parental lines, even though in less abundant levels (between 5% - 25%). These findings together suggest that genetic alterations might not be the major driving force behind EGFR-independent relapse tumor development.

### The relapse is accompanied by glioma subtype reprogramming

The lack of evident genetic alterations in the iEIP relapsed tumors prompted us to question whether epigenetic factors might fuel oncogenic EGFR-independent tumor growth upon dox-induced silencing of EGFR* transgene. To explore this possibility, we next characterized molecular and phenotypic changes along relapse development. As compared to matched treatment-naïve control tumors, western blotting analysis of relapsed iEIP tumors revealed a stark loss of glioma proneural/classical subtype-enriched OLIG2 and ASCL1 expression with a concurrent upregulation of mesenchymal subtype signature genes, VIM and TGFBI (Fig. 2A) (28). IHC analysis further confirmed a ubiquitous silencing of OLIG2 with a concerted VIM upregulation in all relapsed tumors analyzed (Fig. 2B), suggesting a tumor subtype transdifferentiation during the relapse.

**Figure 2.**
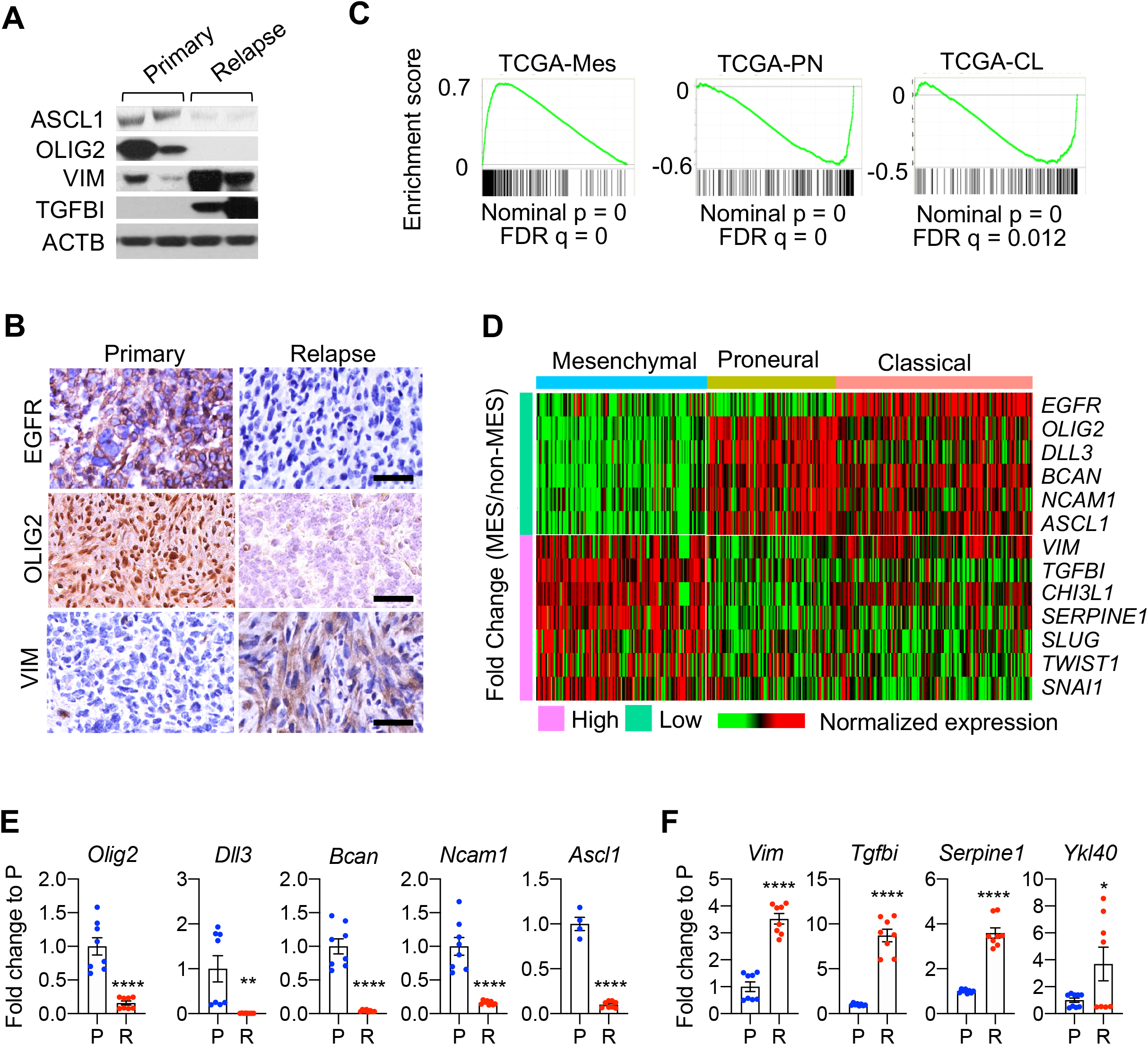
Relapsed tumors following EGFR* inhibition show subtype transition. **A**, Representative images of WB analysis for indicated proteins in primary and recurrent tumors. **B,** Representative images of IHC staining performed on sections of primary and relapsed tumors. Scale bars, 50 μm. **C**, GSEA plots showing positive enrichment of the mesenchymal signature (MES) gene set and negative enrichment of the proneural (PN) and classical signature (CN) gene sets. **D**, Gene expression comparison between mesenchymal (MES) and non-mesenchymal (non-MES) GBMs. The expression profile of 369 IDH wild-type GBMs were compiled from TCGA GBM dataset. Cases were classified into mesenchymal, proneural and classical GBMs according to the latest glioma-intrinsic subtype signatures. Green and pink left bar indicate fold change values (MES vs. non-MES) less and greater than 1, respectively. Heatmap represents the normalized gene expression of thirteen genes, and red and green color indicate high and low expression, respectively. **E-F,** qRT-PCR analysis of primary (P) and relapsed (R) tumors for PN or CL (**E**) or MES (**F**) signature genes in tumor tissues. Mean ± s.e.m. of 4-10 tumors. Statistical significance was determined by unpaired t-test. *P<0.05, **P<0.01, ***P<0.002, and ****P<0.0001.

To identify the molecular components underlying the relapse, we analyzed our previously published expression profiles of three relapsed tumors relative to their paired treatment-naïve controls (27) (GSE64751) to define their molecular subtypes. Using the classifying gene list developed by the Cancer Genome Atlas (TCGA) group (29), gene set enrichment analysis (GSEA) indicated that relapsed tumors were strongly enriched for mesenchymal subtype signature genes, as compared to the treatment-naïve tumors that exhibited features of the classical and proneural subtypes (Fig. 2B and 2C).

Concurrently, gene expression heatmap and further quantitative real-time PCR (qPCR) identified a panel of significantly upregulated mesenchymal-subtype associated genes (*Vim, Tgfbi, Ykl40* and *Serpine1*) as well as a list of silenced proneural- or classical-subtype enriched genes (*Olig2, Dll3, Bcan, Ncam1,* and *Ascl1*) (Fig. 2D-F).

### Enforced mesenchymal transcription factor expression relieves EGFR dependency

In agreement with their mesenchymal fate transdifferentiation, qPCR and western blot analysis of the relapsed tumors revealed a significant upregulation of core mesenchymal transcriptional factors Slug and Twist, as compared to the treatment-naïve controls (Fig. 3A-3C). Importantly, IHC analysis for Slug or Twist indicated that the enhanced expression in the relapsed tumors was pervasive and not confined to a particular pocket area (Fig. 3D). Given that mesenchymal transdifferentiation has been a touted resistance mechanism to anti-EGFR therapies in both clinic and model systems of lung cancer treatment (30–32), our observation raised the possibility that dysregulated mesenchymal transcriptional factor expression might underlie the tumor relapse upon oncogenic EGFR* suppression.

**Figure 3.**
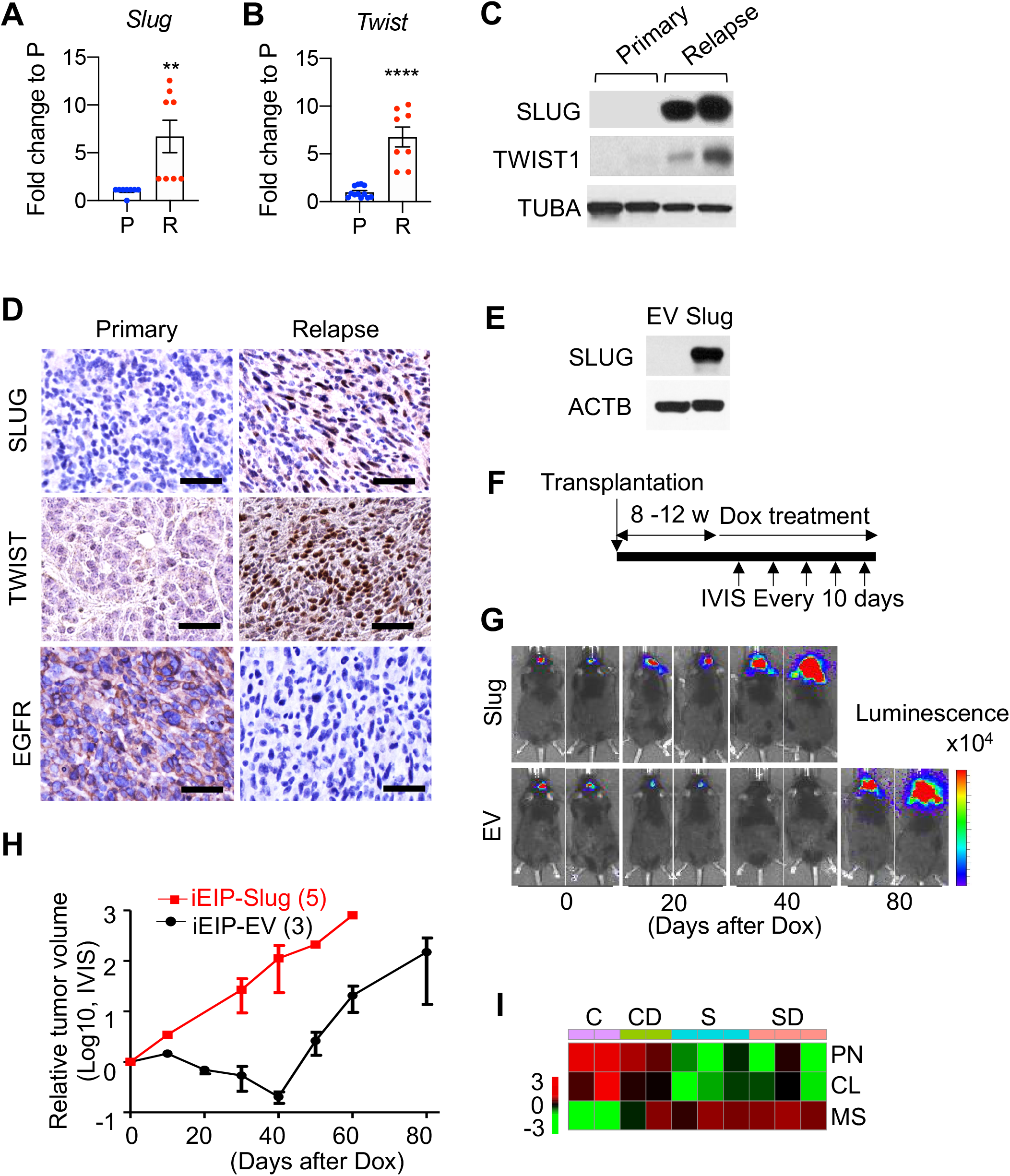
Mesenchymal transcription factor expression-reprogrammed tumors progress independently of EGFR signaling. **A-B,** qRT-PCR analysis of primary (P) and relapsed (R) tumors for Slug (A) or Twist (B). Mean ± s.e.m. of 8 tumors. **C**, Representative images of WB analysis for indicated proteins in primary and relapsed tumors. **D**, Representative images of IHC staining performed on sections of primary and relapsed tumors. Scale bars, 50 μm. **E,** Representative images of WB analysis for SLUG in iEIP cells expressing empty vector (EV) or Slug. **F**, Schema for orthotopic transplantation of control and SLUG iEIP cells, dox treatment and IVIS imaging. **G**, Representative BLI on indicated days post dox treatment. **H**, Tumor growth was measured at indicated times and calculated relative to initial tumor volume. Mean ± s.e.m. of 3-5 biological replicates. Day 0 represents the day when treatment was initiated. **I**, Heat map of C, CD, S and SD showing PN, CL, and MS enrichment. The signature score was calculated using ssGSEA. Statistical significance was determined by unpaired t-test for A. **P<0.01 and ****P<0.0001.

To test this hypothesis, we orthotopically grafted a cohort of immunocompromised *RAG1−/−*recipient mice with luciferase-expressing iEIP glioma cells that were further transduced with either a lentiviral vector control (EV) or Slug expressing construct (Fig. 3E). The animals were kept off-dox and tumor growth was monitored weekly by BLI. After the BLI signals reached ~5 × 10^5^, the animals were switched to dox-containing drinking water to turn off the EGFR* transgene (Fig. 3F). As expected, the EV-transduced control tumors experienced a robust initial response upon dox treatment, followed by an extended period of dormancy before the eventual recurrence following an extended period of dormancy (Fig. 3G and 3H). By contrast, the growth of Slug transduced tumors showed little evidence of response to the dox-induced EGFR* suppression, indicating that upregulation of mesenchymal transcription factors is able to drive oncogenic EGFR-independent iEIP glioma progression. Along this line, further RNA-seq analysis of the four groups of tumor samples (C, control EV-transduced dox-treatment-naïve; S, Slug-transduced dox-treatment-naïve; CD, control EV-transduced relapse under dox-treatment; and SD, Slug-transduced under dox-treatment) confirmed that enforced Slug expression was sufficient to promote mesenchymal transdifferentiation of iEIP tumor cells (Fig. 3I). But notably, despite the fact that enforced Slug expression in iEIP glioma cells overcame their dependency on oncogenic EGFR* signaling, knockdown of Slug expression in relapsed iEIP tumor cells was not sufficient to restore their EGFR* dependency (Supplementary Fig. S2A). Instead, we found a sharply increased Twist1 expression in these Slug-depleted tumors (Supplementary Fig. S2B and S2C), suggesting a functional redundancy among mesenchymal transcriptional factors.

To determine whether mesenchymal transdifferentiation also promoted oncogenic EGFR independency in human gliomas, we firstly transduced patient-derived EGFR^high^ glioma cells with lentivirus encoding either control or SLUG. These cells were then further infected with EGFR-targeting CRISPR/Cas9-based sgRNA (EGFRc) before subcutaneously transplanting into immunocompromised recipient animals (Supplementary Fig. S3A). As expected, the EGFR depleted glioma cells exhibited a significantly diminished tumorigenicity relative to their EV-transduced controls (Supplementary Fig. S3B and 3C). Notably, co-transduction of a SLUG construct completely nullified the tumor suppressive effect of EGFR depletion on tumor progression, although SLUG expression by itself did not appear to have a primary effect on control tumor proliferation rate. These findings are in agreement with the role of mesenchymal core transcriptional factors in initiating glioma fate transdifferentiation and to promote oncogenic EGFR signaling-independent relapse.

### Relapse-inducing cells emerge during the proximal EGFR inhibition response phase

Tumor relapse from targeted therapies often results from a rare subpopulation of treatment-resistant cells that persist through therapy. Indeed, immunofluorescence analysis of dox-treated tumors from orthotopically transplanted iEIP cells identified residues of GFP-positive tumor cells that survived through a 10-day treatment (Fig. 4A). As expected, these potential relapse-inducing tumor cells were depleted of their EGFR* expression. BrdU incorporation analysis indicated that after surviving the initial apoptosis, these residual tumor cells remained relatively non-proliferative or dormant compared to their untreated controls (Fig. 4B). Their non-clustering distribution precludes that these relapse-inducing cells originated from a subclone of tumor cells with pre-existing resistance-conferring mutations.

**Figure 4.**
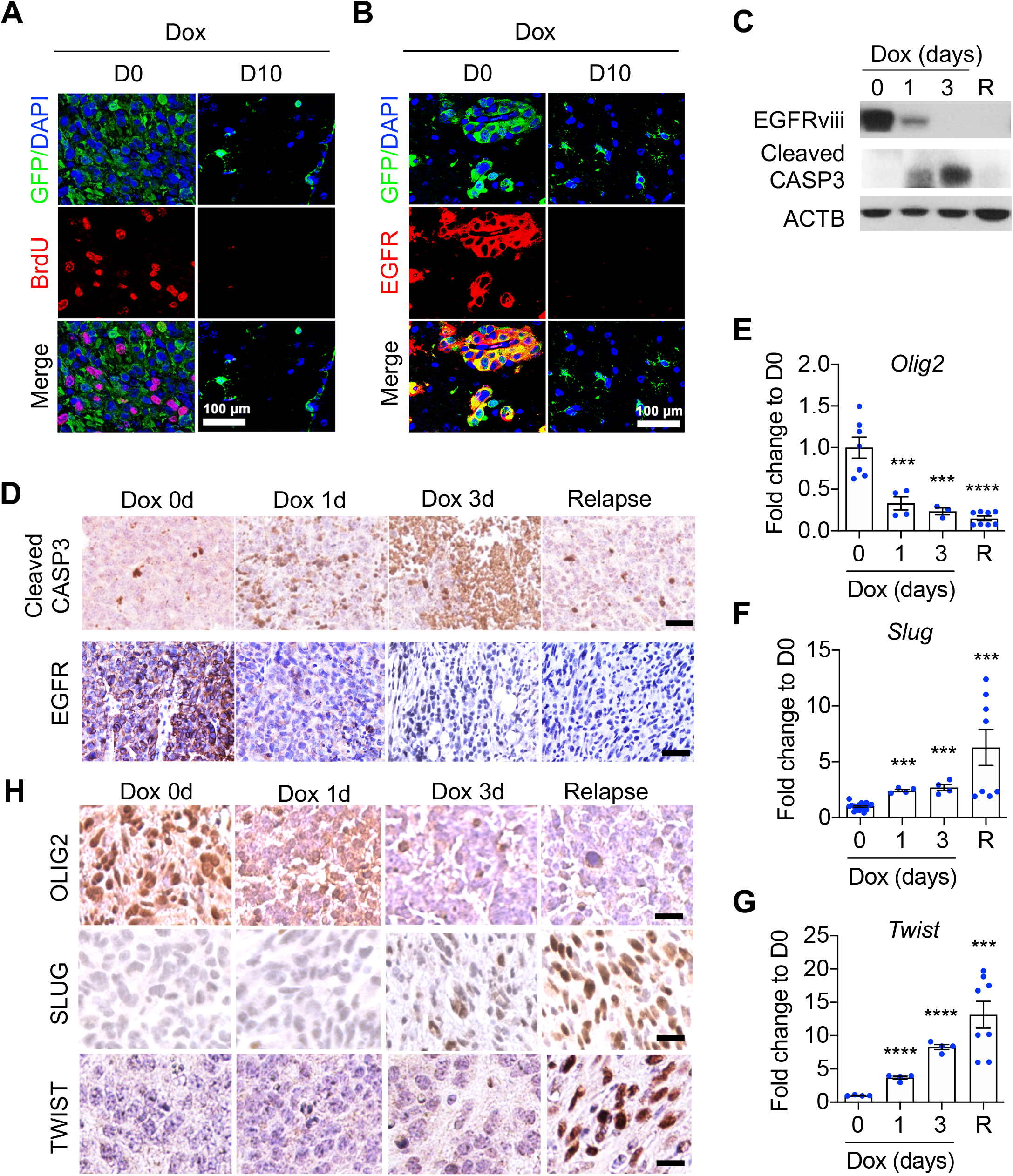
Relapse-inducing cells emerge during the proximal EGFR inhibition response phase. **A-B,** Representative IF analysis for BrdU incorporation (A) and EGFR expression (B) in GFP-labeled iEIP tumors treated with dox for indicated time. Scale bars, 100 μm. **C,** Representative WB analysis for indicated proteins in iEIP tumors. R: relapsed tumor. **D** and **H,** Representative IHC images for indicated proteins in iEIP tumors following dox treatment for indicated time. Scale bars, 50 μm (D), 10 μm (H). **E-G**, qRT-PCR analysis. Data are presented as mean ± s.e.m. of 8 biological replicates. Statistical significance was determined by one-way ANOVA. ***P<0.002 and ****P<0.0001.

To address whether mesenchymal transdifferentiation occurred at the early phase of treatment, we examined acute response of the orthotopically transplanted iEIP tumors to dox-induced EGFR* suppression. Western blot analysis of tumor samples revealed a rapid reduction of transgene-encoded EGFR* protein expression by day 3 of dox treatment (Fig. 4C). Consistent to its role in iEIP tumor maintenance, the prompt reduction of EGFR* expression was accompanied by a rapid increase of cleaved caspase-3 levels, an indicative of apoptosis (Fig.4C and 4D). Along this line, qPCR and IHC analyses further revealed a rapid diminution of proneural marker Olig2 expression upon dox treatment (Fig. 4E). It correlated with a sporadic upsurge of surviving tumor cells expressing mesenchymal transcription factor Slug or Twist1 (Fig. 4F-H). These findings suggest that relapse is likely driven by a stochastic process of mesenchymal transdifferentiation.

### YAP1 activation drives mesenchymal transdifferentiation in response to acute EGFR* suppression

To characterize the transition state following EGFR* deprivation, we performed RNA sequencing (RNA-seq) of orthotopically transplanted iEIP tumors following control or dox treatment. GSEA revealed YAP (Yes-associated protein) signaling (nes=-2, FWER p-value=0) as the top increased gene expression signature of dox-treated tumors (Fig. 5A and 5B). Interestingly, although qPCR assay revealed no consistent Yap1 gene upregulation in response to acute EGFR* suppression (Fig. 5C), IHC analysis of tumor samples following a 1- or 3-day dox treatment revealed a progressively increased nuclear expression of YAP1 (Fig. 5D and 5E). This is consistent with the fact that the regulation of its intracellular localization is a key determinant of YAP activity (33). Notably, nuclear-localized YAP1 was mostly limited to luciferase-expressing tumor cells (Supplementary Fig. S4A).

**Figure 5.**
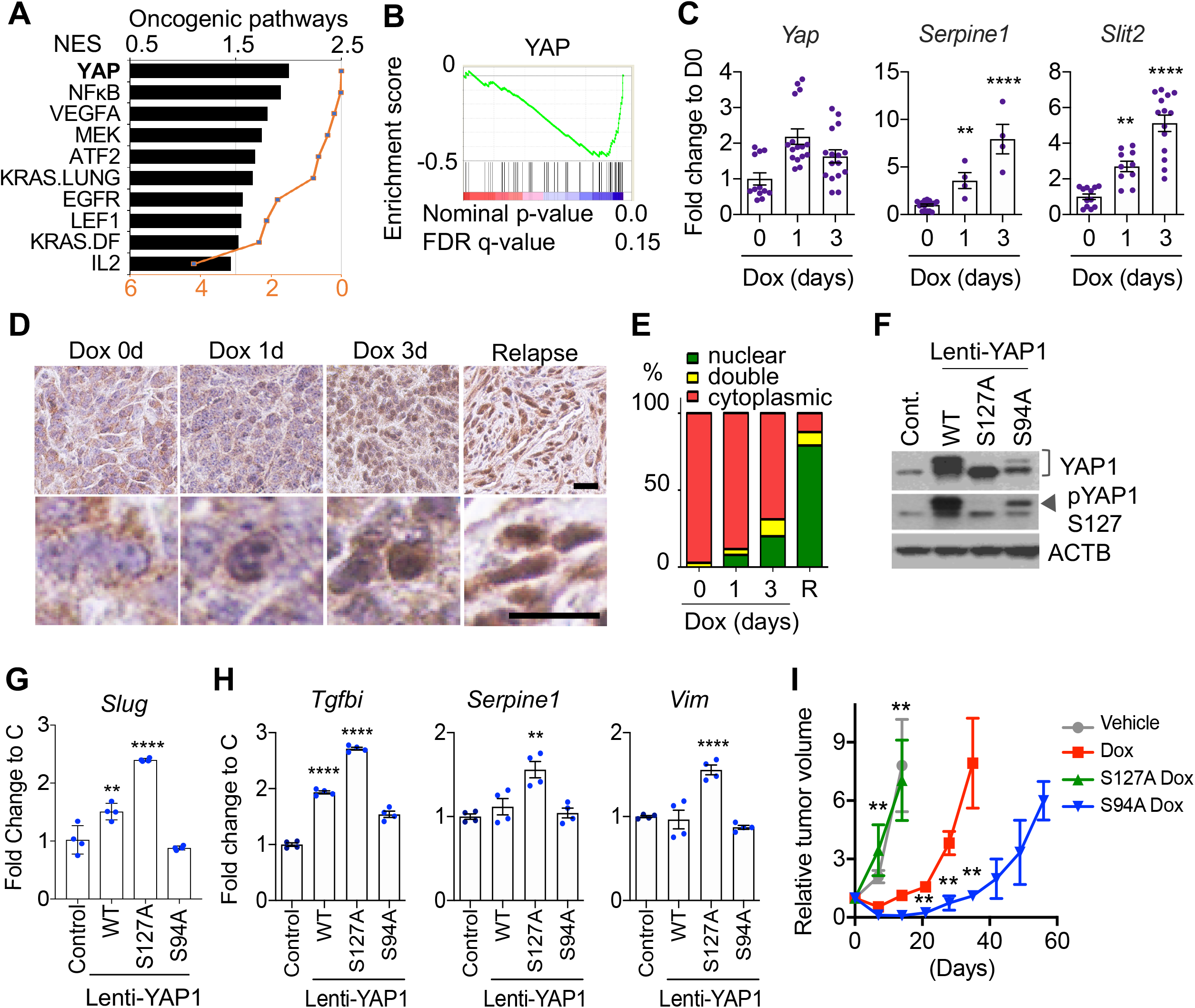
YAP1 activation drives mesenchymal transdifferentiation in response to acute EGFR* suppression. **A**-**B**, GSEA of C vs CD, S, SD tumors. Enriched oncogenic pathways (A). GSEA enrichment plot for YAP (B). **C**, qRT-PCR analysis in iEIP tumors. Mean ± s.e.m. of 4-12 biological replicates. **D,** Representative images of YAP1 IHC. Lower panels are higher magnification from upper ones. Scale bars, 50 μm. **E**, Summary of subcellular localization of YAP1 on control (N=263), dox 1 day (N=212), 3 day (N=210), and relapse tumor (N=168) cells. Results are from four biological replicates. **F**, Representative WB analysis in iEIP cells infected with lenti-virus expressing respective YAP1 constructs. **G** and **H**, qRT-PCR analysis in iEIP cells expressing YAP1 or YAP1 mutant (DA: YAP1^S127A^, DN: YAP1^S94A^). Mean ± s.e.m. of 4 independent experiments. **I,** Relative tumor volume of iEIP subcutaneous tumor. Mice bearing subcutaneously transplanted iEIP cells expressing YAP1^S127A^ (N=6), YAP1^S94A^ (N=6), or control (N=8) were treated with either vehicle or dox containing water. Percent survival of mice sacrificed by the tumor size reached to 200~400 mm^3^. Mean ± s.e.m. of 6-8 tumors. Statistical significance was determined by one-way ANOVA for C, G, H, and I. *P<0.05, **P<0.01, ***P<0.002, and ****P<0.0001.

YAP is the main effector of Hippo signal transduction pathway. Its activation has been implicated in resistance to various targeted therapies, including EGFR tyrosine kinase inhibitors in NSCLC (32,34). To determine whether YAP1 activation contributes to treatment-induced mesenchymal transdifferentiation and relapse development, we transduced primary iEIP glioma cells with retrovirus encoding YAP wild type (YAP^WT^), nuclear-localized active mutant (YAP^S127A^) (35), or mutant (YAP^S94A^) defective of TEAD binding and nuclear translocation (36) (Fig. 5F). qPCR analysis of glioma cells transduced with either YAP^WT^ or active YAP^S127A^, but not YAP^S94A^ mutant, significantly upregulated the expression of Slug and other mesenchymal subtype markers including Tgfbi, Serpine1 and Vim (Fig. 5G and 5H), indicating that YAP activation is capable of driving mesenchymal transdifferentiation.

To determine whether activated YAP could promote oncogenic EGFR* independent glioma growth *in vivo*, primary iEIP cells transduced with vector control or construct encoding YAP^S127A^ or YAP^S94A^ were subcutaneously transplanted into immunocompromised recipient animals. The animals were kept off dox until the tumor reached a palpable size (~ 200 mm^3^). Notably, expression of YAP^S127A^ or YAP^S94A^ did not affect primary tumor growth (Fig. 5I). However, similarly to the Slug-expressing iEIP tumors, the proliferation of active YAP^S127A^ expressing tumors showed little response to dox-induced EGFR* suppression. By contrast, dominant negative YAP^S94A^-expressing tumors showed a much deeper initial regression upon dox-treatment and also took significantly longer to recur, indicating that YAP activity is required for EGFR*-independent relapse.

### Treatment-induced TGFβ secretion activates YAP

Targeted therapy has been shown to induce a network of secreted signals that can instigate drug resistance development by supporting relapse-inducing cells (37–39). To determine whether the anti-EGFR treatment-induced secretion of signaling factors could also promote YAP activation, we generated conditioned media (CM) from control and dox-treated iEIP cultures. Western blot and immunofluorescence analyses revealed that iEIP cells treated with dox-CM had a significantly increased nuclear YAP fraction as compared to the control CM treated samples (Fig. 6A and 6B). These results indicate that EGFR* suppression-stressed iEIP cells respond to secreted YAP-activating factor(s).

**Figure 6.**
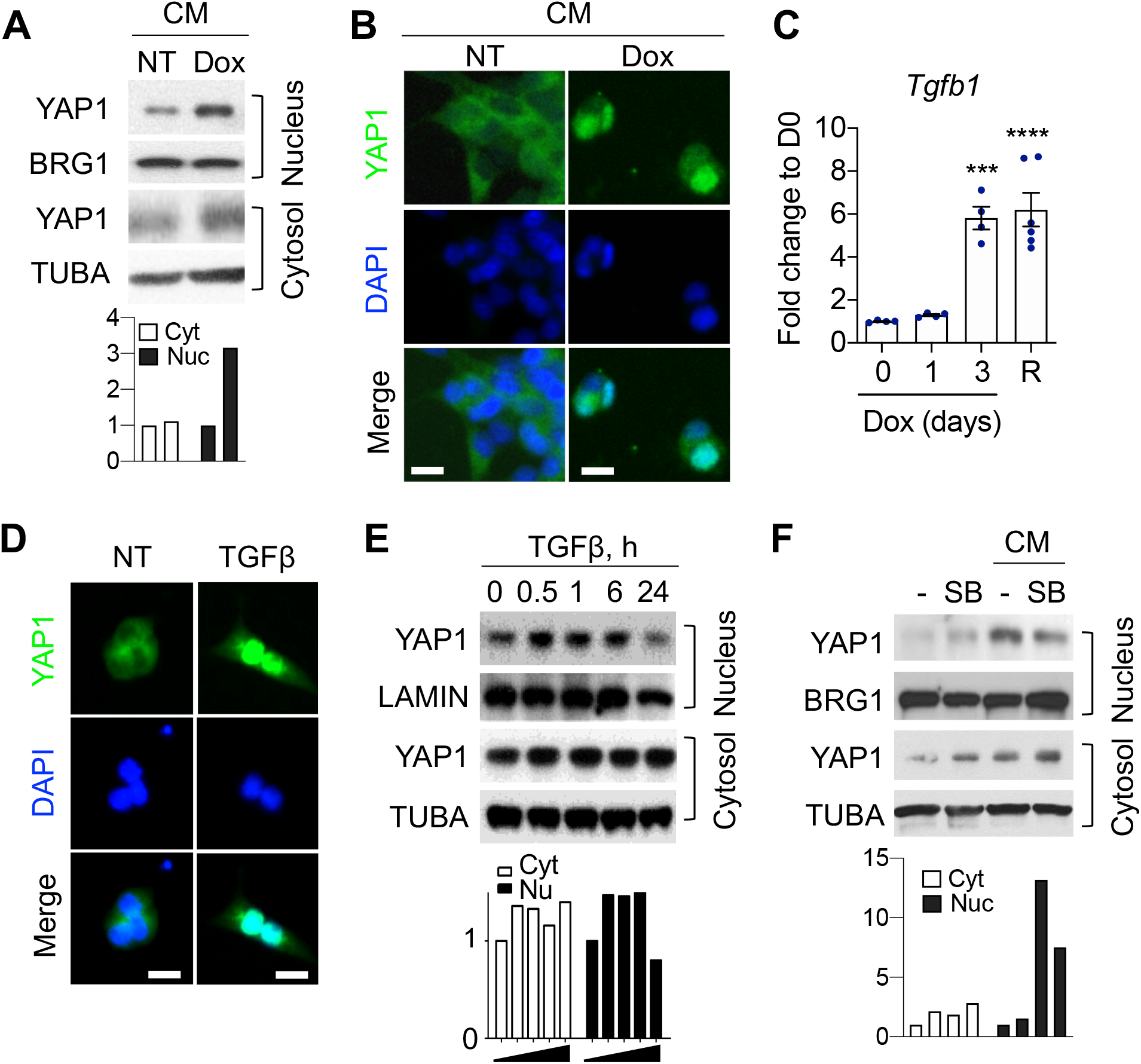
Treatment-induced TGFβ secretion activates YAP. **A-B,** Representative images of WB analysis for nuclear/cytosolic fraction (A) and IF analysis (B) in iEIP cells treated with conditioned medium (CM) of iEIP cells cultured with or without Dox. Scale bars, 10 μm. **C**, qRT-PCR for *Tgfb1* in iEIP tumors. Mean ± s.e.m. of 4-6 tumors. Statistical significance was determined by one-way ANOVA. ***P<0.002 and ****P<0.0001. **D-E,** Representative images of IF analysis (**D**) and WB analysis for nuclear/cytosolic fraction (**E**) in iEIP cells treated with 10 ng/ml of TGFβ1. Scale bars, 10 μm. **F**, Representative images of WB analysis for nuclear/cytosolic fraction in iEIP cells treated with conditioned medium (CM) of iEIP cells cultured with or without 10 μM of SB505124. The bar graphs under WB images of A, E, and F are quantification for each blot.

To identify the relevant YAP-activating factor(s), we next analyzed gene expression changes and identified a panel of cytokines (i.e. GRN, FIGF, TNFA, SLIT2, and TGFB1) whose expression was significantly augmented in relapsed iEIP tumors compared to their treatment-naïve controls (Supplementary Fig. S5A and S5B). To determine their role in YAP1 activation and mesenchymal transdifferentiation during EGFR* suppression, we treated the cultured primary iEIP cells with the factors individually and assessed their mesenchymal transdifferentiation potential. qPCR analysis revealed that treatment of TNFα, GRN, FIGF, or SLIT2 minimally affected mesenchymal subtype gene expression (Supplementary Fig. S5C-S5E). In contrast, TGFβ stimulation significantly augmented Serpine1, Tgfbi and Vimentin mRNA transcription. In addition, TGFβ expression in iEIP tumors was greatly elevated in the acute phase of EGFR* inhibition, tracking with the timeline of YAP1 activation (Fig. 6C). Concordantly, immunofluorescence and western blot of subcellular fractionates of cultured iEIP cells revealed a strong increase of YAP1 nuclear translocation upon TGFβ treatment (Fig. 6D and 6E). Conversely, inhibition of TGFβ signaling by TGFβR1 inhibitor SB525334 compromised dox-CM-induced YAP nuclear accumulation (Fig. 6F). Together, these findings identify the TGFβ/YAP/Slug signaling axis as a crucial mediator of treatment-induced mesenchymal reprograming and anti-EGFR resistance development.

### Activation of YAP in recurrent gliomas predicts poor patient survival

Having established that YAP activation stimulates mesenchymal transdifferentiation and relapse from treatment, we next questioned whether YAP activation also correlated with mesenchymal transition in recurrent clinical samples. We derived a panel of total twenty-two YAP core signature genes from the molecular signature database (MSigDB) that were also significantly up-regulated in EGFR*-independent iEIP tumors (CD, S, and SD) as compared to treatment-naïve tumors (C) (Fig. 7A). By analyzing gene expression profiles from our previously published cohort of 91 recurrent GBM patient samples (28), 39 cases were classified as mesenchymal subtype (MES), while the rest 52 as non-mesenchymal subtype (non-MES). Wilcoxon rank-sum test revealed a significant increase of the signature score for YAP core target genes (ssGSEA) as well as YAP1 expression in the MES subtype relative to the non-MES group (Fig. 7B). Consistently, IHC analysis in 22 recurrent GBM samples revealed a strong correlation between nuclear-localized YAP1 levels and core mesenchymal transcriptional factor SLUG protein expression (p<0.0001, R^2^=0.78, Fig. 7C and 7D).

**Figure 7.**
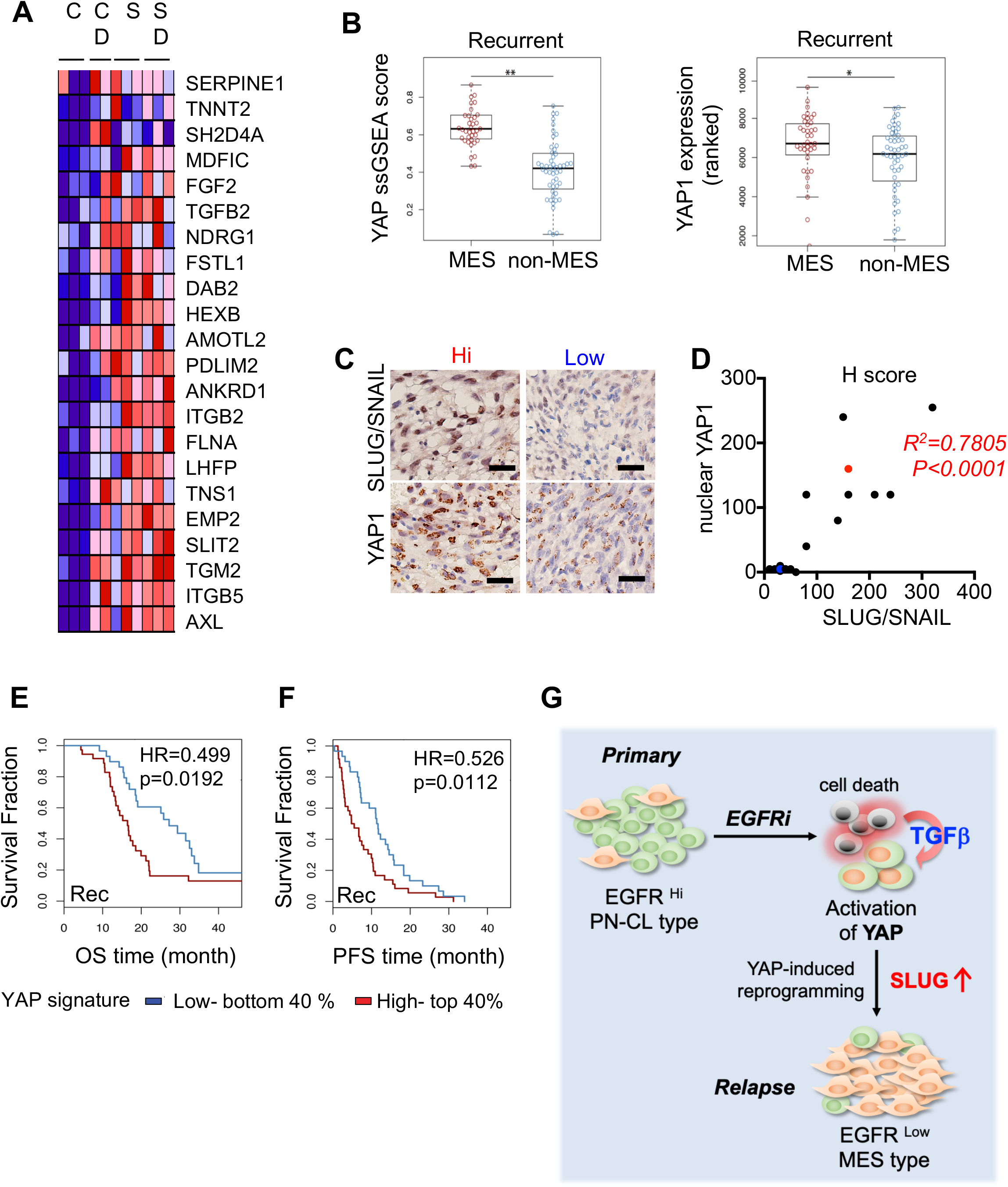
Activation of YAP in recurrent gliomas predicts poor patient survival. **A,** Heat map of 22 YAP core signature genes significantly up-regulated in EGFR*-independent tumors. **B,** The comparison of the average score for YAP pathway or expression of YAP1 between MES and non-MES in recurrent tumors. *P<0.05 and **P<0.01 **C-D,** Recurrent GBM patient samples (N=22) were scored for YAP1 and SLUG/SNAIL nuclear expression. Representative images of YAP1 IHC (**C**) and H-score plot (**D**). **E-F,** Survival analysis of paired IDH1 WT GBM patients (N=91). Overall survival (**E**) and PFS (**F**) analyses of recurrent patients with either high or low YAP core signature expressions. **G,** Schema for mechanism of TGFβ-YAP1 activation-SLUG induced mesenchymal transition under EGFR targeted therapy.

Lastly, we evaluated the effect of YAP activation on patient survival in 54 cases for whom annotation on overall survival time (OS) and time to disease progression (progression free survival, PFS) were available. Patients whose recurrent tumors were classified as high YAP-ssGSEA (top 40%) trended toward adverse overall survival (log rank test p = 0.0192 with HR = 0.499) (Fig. 7E) and PFS (log rank test p = 0.0112 with HR = 0.526) (Fig. 7F). Collectively, these results together suggest that YAP-dependent mesenchymal program represents a highly relevant molecular pathway that determines disease recurrence, progression and prognosis.

## Discussion

Targeted-therapy is the standard care for many cancer types that harbor an activated oncogene. But even with effective therapies that lead to partial or complete response, it is expected that rare tumor cells always survive and eventually re-initiate the malignant disease with dismal outcome (40–42). In this study, we applied an inducible oncogenic EGFR-driven glioma mouse model to investigate anti-EGFR therapeutic response and resistance development. By analyzing the proximal and distal response upon oncogenic EGFR* deprivation, our study identified treatment-induced cell plasticity as an underlying mechanism that drives EGFR-targeted therapy evasion and relapse development. Our findings revealed that anti-EGFR therapy activated a TGFβ/YAP/Slug signaling cascade that subsequently instigated mesenchymal lineage transdifferentiation in a rare population of glioma relapse initiating cells. We further demonstrated that this process enabled reprogramming of the oncogenic EGFR-addicted glioma cells toward a phenotypic state no longer relying on EGFR signaling (Fig. 7G). Inhibition of this adaptive response through enforced dominant negative YAP expression significantly delayed anti-EGFR relapse from anti-EGFR treatment and prolonged animal survival.

EGFR is amplified and mutated in a majority of human malignant gliomas; yet various treatment strategies targeting EGFR have thus far failed in clinical trials (15). One major reason might be insufficient target efficacy due to either suboptimal mutant EGFR inhibition (16,17), or inefficient drug penetration and distribution in the CNS (43,44). Additionally, a body of reports also pointed that intratumoral heterogeneity or compensatory upregulation of other RTKs, including PDGFRA and MET, could contribute to anti-EGFR therapy resistance (18–20,45). Similarly, activation of downstream effectors or engagement of alternative survival pathways could also mediate EGFR targeted therapy evasion (21,22,46). In addition to aforementioned anti-EGFR resistance mechanisms that rely on activation of compensatory signaling effectors of the same or alternative survival pathways, our study points to the importance of glioma cell plasticity as another avenue of therapy evasion. Indeed, even with near complete oncogenic EGFR deprivation that leads to deep response, we found that a small population of iEIP glioma cells would survive and undergo treatment-induced transdifferentiation towards a mesenchymal-like state that no longer depends on oncogenic EGFR*. These findings indicate that anti-EGFR therapeutic resistance in malignant glioma might result, at least in part, from treatment-induced cell plasticity and enhancement of mesenchymal transdifferentiation.

The mesenchymal switching, an important biological process that enables cells to reprogram towards distinct phenotypes in response to their environmental changes, is identified not only during development such as mesoderm and neural tube formation, but also upon injury and disease (47,48). In cancers, epithelia to mesenchymal transition (EMT) has been widely associated with tumor stemness, metastasis and drug resistance (42,49). The development of EMT as a drug-resistant mechanism has been observed in various in vitro and in vivo model systems of epithelia cancers, including EGFR mutant lung cancer (31,32), KRAS mutant colon and pancreatic cancers (50,51), and PyMT-driven breast cancer (52). In malignant gliomas, transcriptome comparison of paired primary and recurrent GBM samples following standard therapy also revealed a trend of phenotypic and molecular shift towards a mesenchymal state during recurrence (53), suggesting that mesenchymal tumor cells may intrinsically lack sensitivity to many conventional and targeted therapies. Besides mesenchymal transformation, other forms of lineage plasticity such as drug-induced neuroendocrine lineage transdifferentiation, has also been implicated in therapeutic resistance development in NSCL and prostate cancer (30,54–56), further underscoring cell plasticity as a general resistance mechanism against conventional and targeted therapies.

Our study reveals that anti-EGFR treatment induces glioma mesenchymal lineage transdifferentiation and oncogenic EGFR-independent relapse through activation TGFβ/YAP/Slug signaling axis. TGFβ, a well-known EMT inducer, has been associated with reduced treatment effectiveness in many types of cancers (57–59). Our findings indicate that therapy-triggered TGFβ secretion promotes YAP nuclear shuffling and subsequent upregulation of mesenchymal transcriptional factor Slug. This is consistent with recent reports that causally link the mesenchymal transformation with YAP-mediated bypass of EGFR or KRAS inhibition (32,50). Similarly, we observed that while inhibition of YAP signaling by itself had little effect on primary iEIP glioma progression, combinatorial suppression of YAP and EGFR* signaling dampened the anti-EGFR treatment-induced mesenchymal transdifferentiation and delayed relapse development. Together, these indicate that activation of YAP signaling can drive mesenchymal transdifferentiation to promote therapy evasion in cancer treatment.

Relapse under targeted therapy often results from rare treatment-refractory tumor cells that persist through treatment (40,41). In this study, we identified a small portion of Slug and/or Twist-positive iEIP glioma cells that could survive initial EGFR* deprivation-induced apoptosis and potentiate later tumor regrowth. Interestingly, those mesenchymal-like relapse-initiating cells do not seem to have pre-existed in treatment-naïve tumors but rather have arisen stochastically following the treatment. These findings therefore support the facultative transient resistance model in which a small portion of tumor cells transiently acquire drug-refractory state by epigenetic modification (42). One possible scenario is that primary iEIP glioma cells may dynamically express fluctuating levels of mesenchymal transcription factors such as Slug and/or Twist. Only cells that display high levels of expression at the time of treatment can survive and assume a transient drug-refractory state. And the final relapse would thus rely on establishment of a stable anti-EGFR resistant state, a process that is presumably enabled by TGFβ/YAP/Slug signaling-mediated mesenchymal transdifferentiation under treatment. This targeted therapy-induced response is likely to be of general relevance to oncogene-driven cancers. Thus, mechanism-based blocking of lineage transdifferentiation may aid the direction of future therapeutic approaches for malignant gliomas.

## Materials and Methods

### Mice

All mouse strains were housed in a barrier facility under protocols approved by the Institutional Animal Care and Use Committee of Weill Cornell Medicine. Rag^−/−^ mice were purchased from the Jackson Laboratories (JAX stock #002216), bred at the institution’s animal facility, and used for graft or xenograft between 8 and 12 weeks of age.

### Cell culture

Sphere culture of iEIP cells was established and maintained in serum-free DMEM/F12 medium (Sigma), containing ITS (Invitrogen), epidermal growth factor (EGF) (20 ng/ml, Peprotech), and basic fibroblast growth factor (bFGF) (20 ng/ml; Peprotech) as previously described (60). The human glioma initiating cells (GS7-11, GS8-11, GSC280) were maintained in serum-free DMEM/F12 medium (Sigma), containing B27 (Invitrogen), 20 ng/ml EGF, and 20 ng/ml bFGF.

### Grafting and xenografting

Orthotopic and subcutaneous grafting experiments were performed as described previously (27). In brief, low passage iEIP glioma cells were retrovirally transduced with mCherry/luciferase expression cassette. 8- to 12-week-old RAG1^−/−^ mice were anesthetized and restrained using a stereotaxic instrument (Stoelting). 50,000 iEIP glioma cells in 2 μl PBS were injected using a Hamilton syringe into right caudate nucleus 2.2 mm below the surface of the brain though a hole made by a dental drill on skull 1 mm anterior and 1 mm lateral from the bregma. For subcutaneous grafting, iEIP glioma cells were resuspended in 50% Matrigel (BD Bioscience; #356231) in PBS and ~10,000,000 cells were injected into each flank of RAG1^−/−^ mice. Tumor growth was monitored and measured every 7 days by caliper, and volume was calculated by the formula: 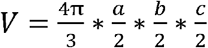 (V tumor volume; a tumor length; b tumor width; c tumor height). Relative tumor volume changes were calculated by dividing by the tumor volume at the beginning of treatment.

### *In vivo* bioluminescent imaging (BLI)

The RAG1^−/−^ mice that were engrafted with iEIP tumor cells expressing the luciferase reporter, were i.p. injected with D-Luciferin (150 mg/kg, Goldbio). Bioluminescent signal was measured using the Xenogen IVIS Lumina system (Caliper LS, Hopkinton, MA). Data acquisition and analysis was performed using the Living Image® software (Caliper LS). All the mice were imaged between 1 to 5 min after the luciferin injection.

### Production of viral particles

pLKO.1-shmSlug4 was a gift from Bob Weinberg (Addgene plasmid # 40648), pPGS-hSLUG.fl.flag was a gift from Eric Fearon (Addgene plasmid # 25696). pHAGE vectors for YAP^WT^, YAP^S127A^, YAP^S94A^ and pLKO-shYap1 were prepared as previously described (61). To generate lentivirus, 1.5 × 10^7^ 293T cells in 150Lmm tissue culture dishes were transfected with 18☐μg of each plasmid DNA along with 4.5 μg of pMD2.G (Addgene #12259) and 9 μg of psPAX2 packaging vectors (Addgene #12260) using polyethylenimine (Polysciences, Warrington, PA, USA; #23966-2). The medium containing lentiviral particles were collected at 48 and 72 h after transfection. To precipitate viral particles, the medium was transferred to an ultracentrifuge tube and spun at 23,000 rpm for 1.5 h at 4☐°C in a Beckman SW28 swinging bucket rotor. Viral particles were resuspended in Opti-MEM (Gibco) on shaker overnight.

### Immunoblots

Total lysates of tumor tissues and cultured cells were prepared in lysis buffer (150 mM NaCl, 50 mM Tris pH 7.4, 0.5% Sodium deoxycholate, 0.1% SDS, 1% NP40), and were cleared by centrifugation. Protein concentrations were quantified with the Pierce BCA Protein Assay Kit (Thermo). The lysates, for histone extraction, were sonicated with pellet 5 times at 20 kHz for 5 second. For Western blotting, 20 μg of protein extract per sample was loaded on acrylamide gel and transferred to PVDF membrane (Millipore). All lysis buffers contained a cocktail of protease inhibitors and phosphatase inhibitors. Antibody sources are described in Supplementary Table I. Secondary antibody, anti-rabbit- or anti-mouse-HRP (Thermo; 1:10,000 dilution), was incubated for 1 h at room temperature. After washing, chemiluminescence was visualized with SuperSignal West Pico (Thermo).

### Immunohistochemistry

Mouse brains or subcutaneously grafted tumors were fixed with 10% (w/v) neutral-buffered formalin, embedded in paraffin, sectioned into 5 μm, deparaffinized, and rehydrated. Antigen epitopes were retrieved by heating in citrate buffer. Primary antibody sources are described in Supplementary Table I. After primary antibody incubation and PBS washes, sections were incubated for 1 h with biotinylated secondary antibodies, followed by applying the ABC kit (Vector labs) and the peroxidase/diaminobenzidine (DAB) method to visualize signals under light microscopy, or incubated with fluorophore-conjugated secondary antibodies (Thermo) to visualize signals under fluorescence microscopy.

### Quantitative real-time RT-PCR

Total RNAs were isolated from tissues and cells using TRIzol Reagent (Thermo) and treated with DNase I (Sigma). cDNAs were prepared with Maxima® first strand cDNA synthesis kit for RT-PCR (Thermo). Quantitative RT-PCR was performed using the Power SYBR® Green Master mix (Applied Biosystems) on 7500 Fast Real-Time PCR System (Applied Biosystems). Values were normalized to 18s rRNA expression. All PCR primers are summarized in Supplemental Table II.

### RNA-seq and data analysis

Total RNAs were isolated and purified from the tumor tissues and subjected to RNA sequencing at the Genomics Resources Core facility of Weill Cornell Medicine. RNA-seq libraries were prepared using the Illumina RNA-seq Preparation Kit and sequenced on Illumina HiSeq 4000 sequencer. RNA-seq reads were mapped using TopHat with default settings. TopHat output data was then analyzed by Cufflinks to calculate FPKM values for known transcripts in the mouse genome reference and variance was estimated to calculate the significance of observed changes in expression. Data is available as GSE108658.

For GBM molecular subtype analysis and primary-recurrent paired data analysis the datasets from (62) was used. Analysis was performed as previously described as in (28). Briefly, U133A array profiles for 533 primary GBM were obtained from the TCGA portal https://tcga-data.nci.nih.gov/tcga/. Mutation calls and DNA methylation profiles were obtained for all samples, where available. GBMs were identified as IDH wild-type while both mutation calls on IDH1 genes were wild-type and GCIMP status inferred using DNA methylation profile was negative. A set of 369 TCGA GBMs were identified as IDH wild-type according to this procedure. Processed primary/recurrence expression data could be analyzed through GlioVis portal http://recur.bioinfo.cnio.es/ (62), which includes 124 primary/recurrent pairs of gliomas and 91 pairs identified as IDH wild-type GBM pairs.

The YAP core signature genes were obtained from the molecular signature database (MSigDB) (http://software.broadinstitute.org/gsea/index.jsp), filtered by in-house iEIP tumor expression profiles, and consequently contained: SERPINE1, TNNT2, SH2D4A, MDFIC, FGF2, TGFB2, NDRG1, FSTL1, DAB2, HEXB, AMOTL2, PDLIM2, ANKRD1, ITGB2, FLNA, LHFP, TNS1, EMP2, SLIT2, TGM2, ITGB5, and AXL. The signature score of YAP core genes was then calculated using ssGSEA (62).

Gene Set Enrichment Analysis (GSEA) was used to identify altered cellular processes and oncogenic pathways of primary and recurrent tumors based on gene sets by Verhaak (29). GESA was performed using c5 and c6 libraries from version 6.0 of MSigDB.

### Whole Exome Sequencing

We performed exome sequencing on three cultured samples from replased tumors (147, 175, relapse 8) as well as two untreated glioma samples (QX, 2QX). Data analysis was carried out using BGI’s proprietary standard pipeline to produce SNP/mutation and InDel calls. To identify possible common mutations arose in replased tumors, we further filtered off low quality mutation calls by increasing the threshold of QD to 10 and DP to 50. Identified loci were visually confirmed by IGV viewer, Data is available as GSE108658.

### Statistical analysis

We determined experimental sample sizes on the basis of preliminary data. All results are expressed as mean ± s.e.m. GraphPad Prism software (version 7, San Diego, CA) was used for all statistical analysis. Normal distribution of the sample sets was determined before applying unpaired Student’s two-tailed t-test for two group comparisons. One-Way ANOVA was used to assess the differences between multiple groups. The mean values of each group were compared by the Bonferroni’s post-hoc procedure. Differences were considered significant when P<0.05.

## Acknowledgements

We thank Dr. Jan Koster at the AMC for support with data analysis and Ms. A. Biswas for proofreading the manuscript.

## Abbreviations

EGFR: epidermal growth factor receptor
GBM: glioblastomas
YAP1: yes associated protein 1
TGFβ: Transforming growth factor beta
TKI: tyrosine kinase inhibitors
PTEN: phosphatase and tensin homolog
GFAP: glial fibrillary acidic protein
BLI: bioluminescence imaging
GSEA: Gene Set Enrichment Analysis

