## Supplemental Figure 1-5 and table S1-2 for "Therapy-induced transdifferentiation promotes glioma growth independent of EGFR signaling"

**Supplementary Figure 1.**

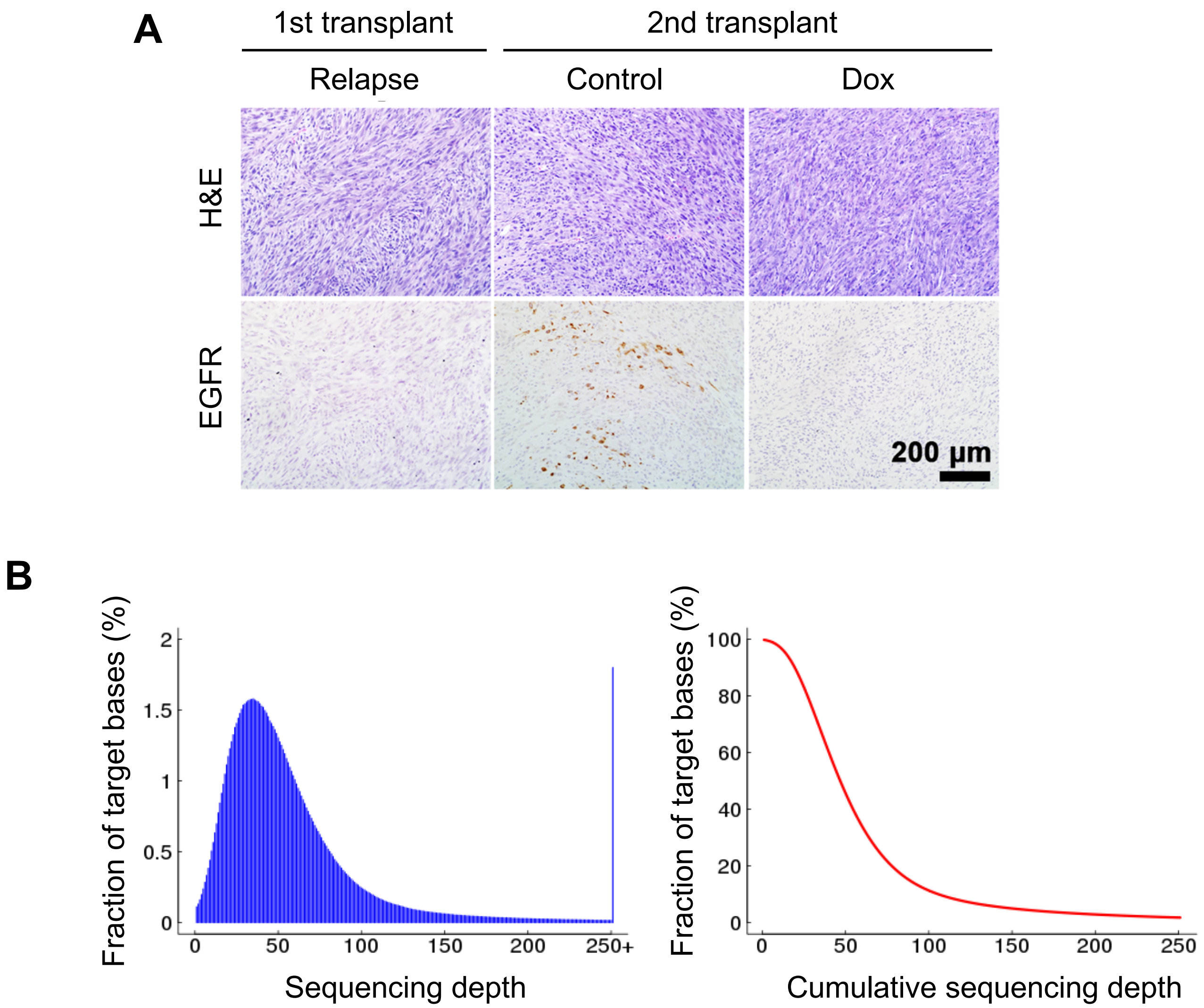


**Supplementary Figure 1. A,** Representative images of H&E and IHC staining against EGFR. Scale bar, 200 µm. **B,** The distribution of per-base sequencing depth and cumulative depth distribution in target regions for a representative sample.

**Supplementary Figure 2.**

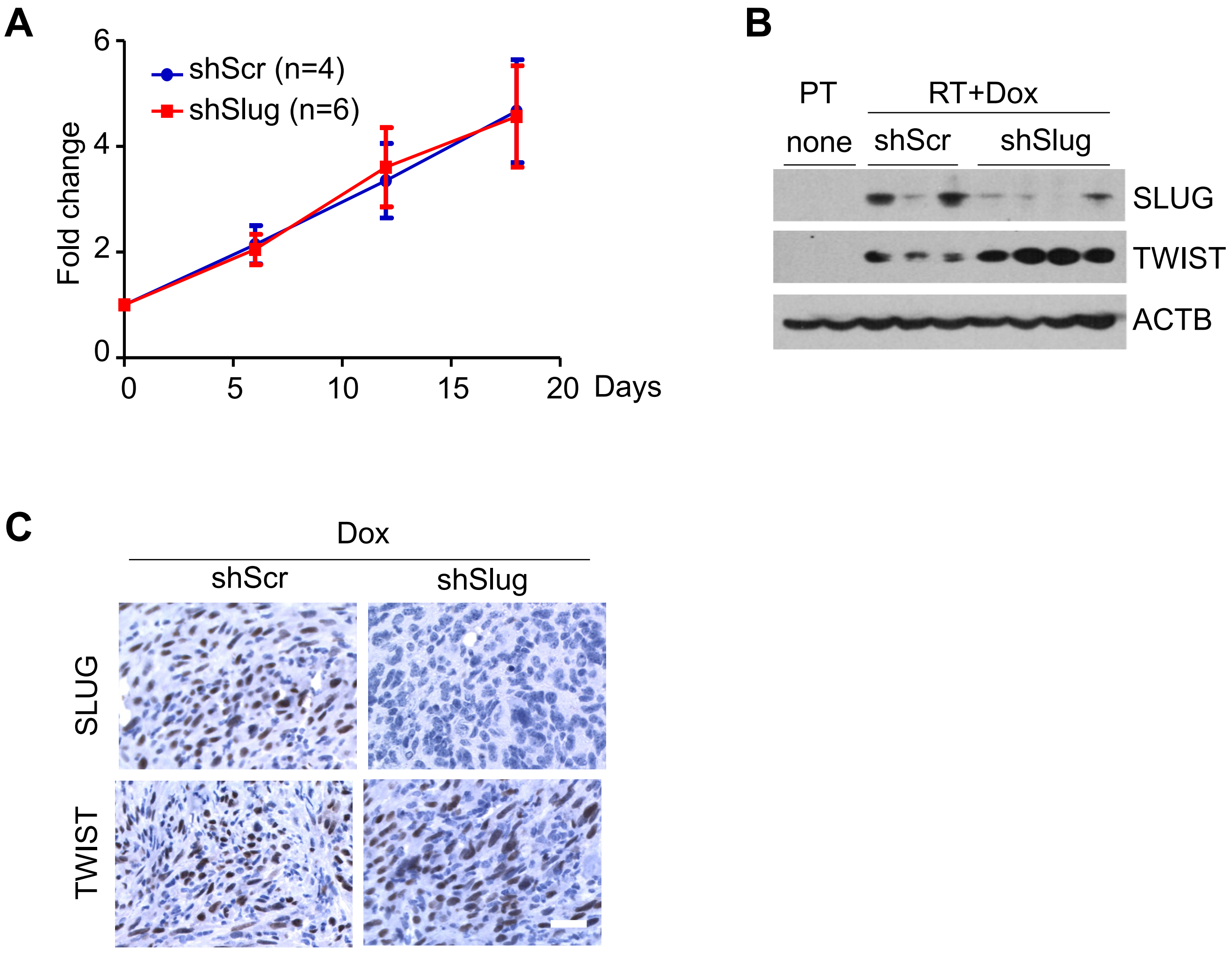


**Supplementary Figure 2. A,** Tumor growth was measured at indicated time and calculated relative to the initial tumor volume. Data, mean ±s.e.m. Day 0 represents the day when tumor was measurable. **B**, WB of primary tumor (PT) and shScr, shSlug infected relapse tumor (RT) treated with doxycycline. **C,** Representative images of IHC staining against SLUG and TWIST performed on sections from shScr and shSlug treated with doxycycline. Scale bar, 50 μm.

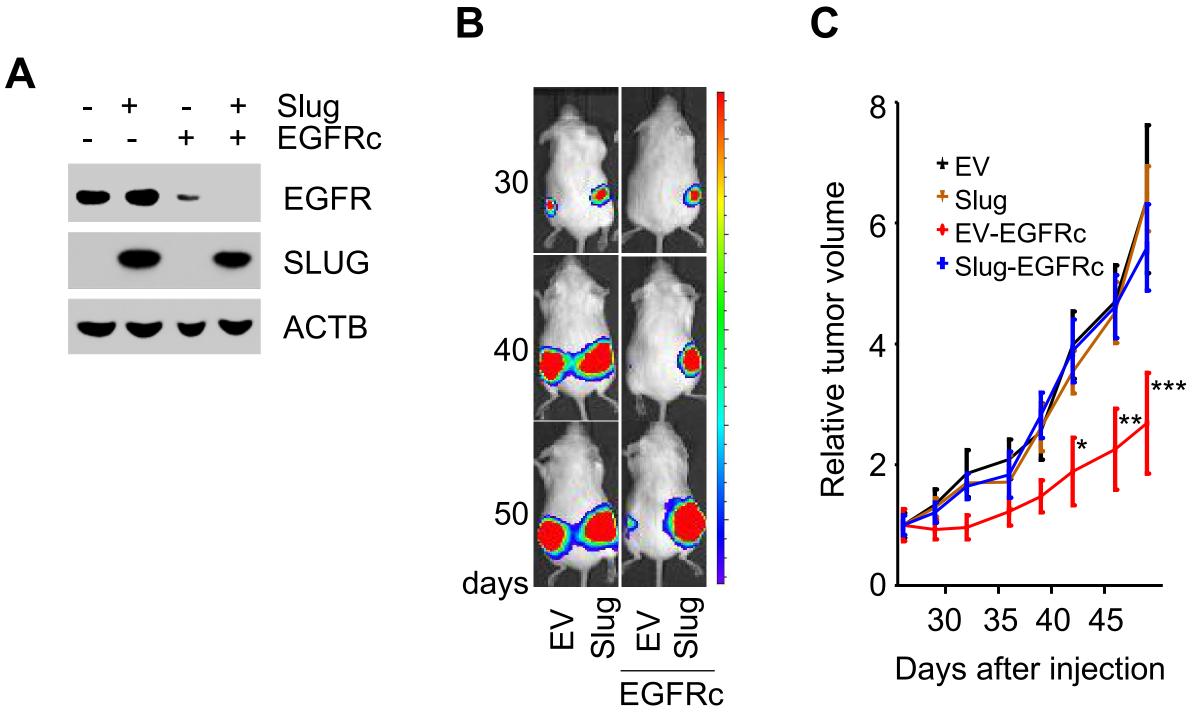


**Supplementary Figure 3. A,** WB for indicated proteins from iEIP cells infected with Slug or sgRNA for EGFR (EGFRc). EV, empty vector. **B,** Representative BLI at indicated time after transplantation of iEIP cells expressing Slug or EGFRc. **C,** Tumor growth was measured at indicated time and calculated relative to the initial tumor volume. Data, mean ± s.e.m. of 3 biological replicates. Statistical significance was determined by one-way ANOVA. *P<0.05, **P<0.01, and ***P<0.001.

**Supplementary Figure 4.**

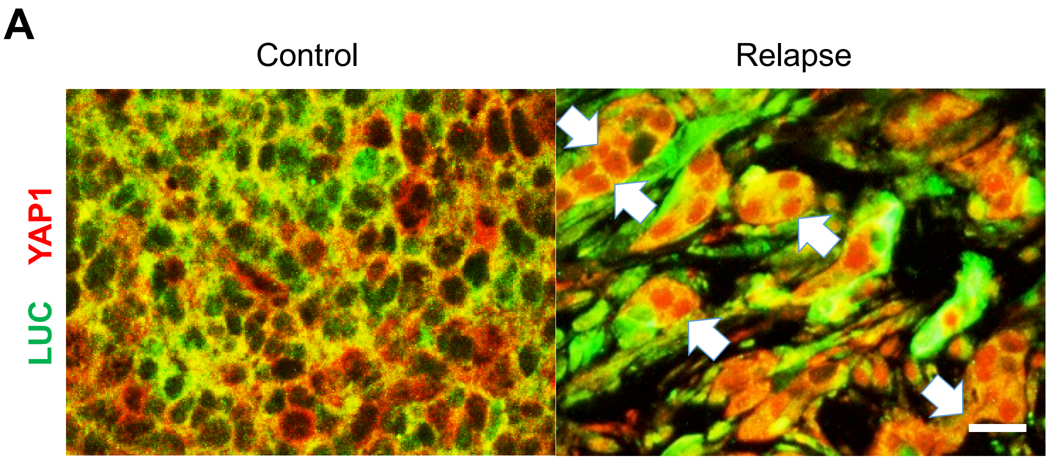


**Supplementary Figure 4. A,** Confocal microscope images co-IF staining for Luciferase (green) and YAP (red). Arrows point to luciferase expressing tumor cells with nuclear YAP1. Scale bar, 50 µm.

**Supplementary Figure 5.**

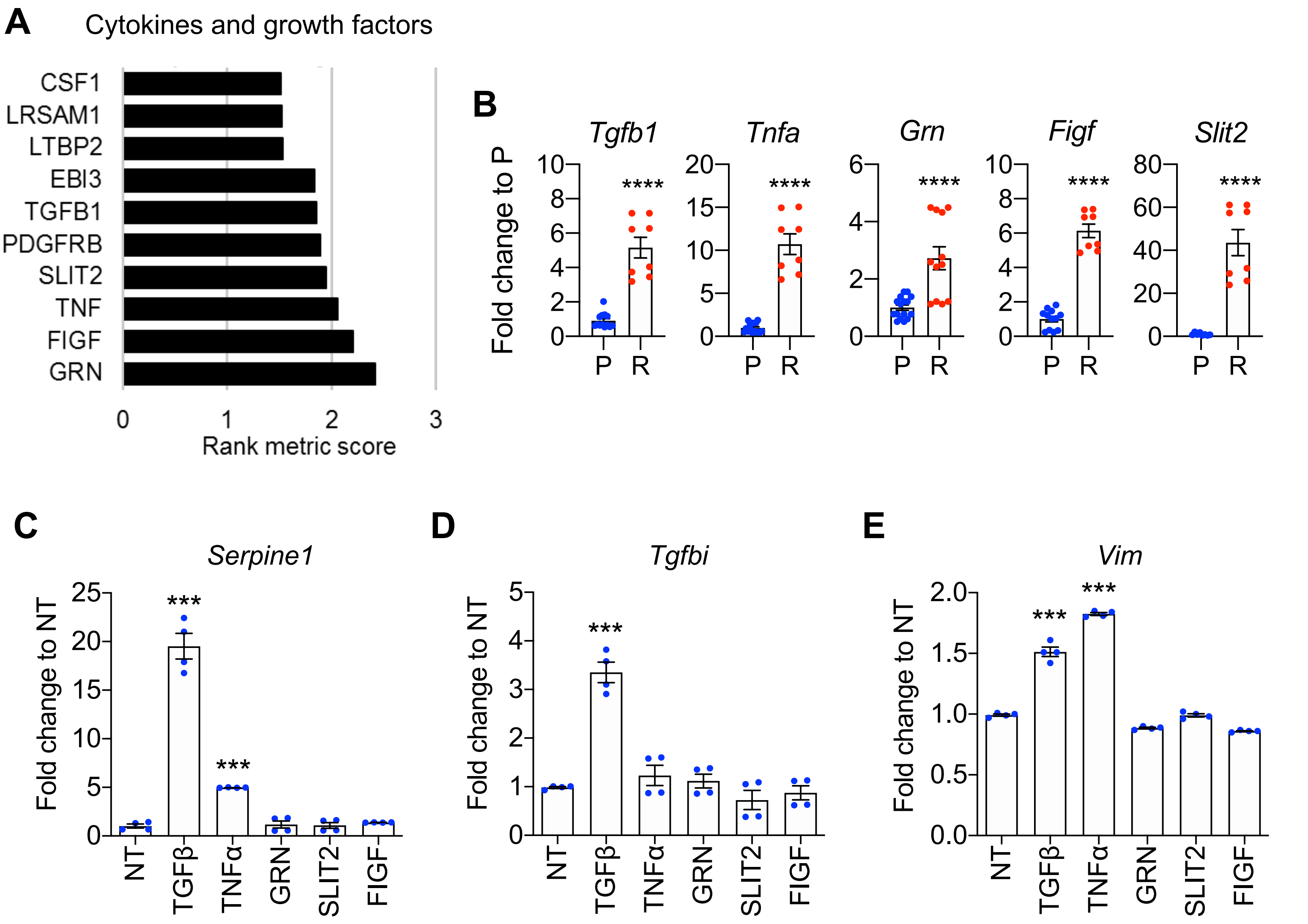


**Supplementary Figure 5. A,** Enriched cytokines and grow factors. **B,** qRT-PCR of mRNA expression of up-regulated cytokines in EGFRviii independent tumor. Mean ± s.e.m. of 8 tumors. **C-E,** mRNA expression of MES-type marker genes *Serpine1* (C), *Tgfbi* (D) and *Vim (*E) in cytokines treated iEIP cells. Gene expression was normalized relative to 18S. Mean ± s.e.m. of 4 independent experiments. Statistical significance was determined by unpaired t-test for (B) and ANOVA for (C-E). **P*<0.05, ***P*<0.01,***P<0.002, and *****P*<0.0001.
